# Gonadectomy and blood sampling procedures in small size teleost models

**DOI:** 10.1101/2020.08.30.271478

**Authors:** Muhammad Rahmad Royan, Shinji Kanda, Daichi Kayo, Weiyi Song, Wei Ge, Finn-Arne Weltzien, Romain Fontaine

## Abstract

Sex steroids, produced by the gonads, play an essential role in the neuroendocrine control of reproduction in all vertebrates by providing feedback to the brain and pituitary. Sex steroids also play an important role in tissue plasticity by regulating cell proliferation in several tissues including the brain and the pituitary. Therefore, investigating the role of sex steroids and mechanisms by which they act is crucial to better understand both feedback mechanism and tissue plasticity. Teleost fish, which possess a higher degree of tissue plasticity and variations in reproduction strategies compared to mammals, appear to be useful models to investigate these questions. The removal of the main source of sex steroid production using gonadectomy together with blood sampling to measure steroid levels, have been well-established and fairly feasible in bigger fish and are powerful techniques to investigate the role and effects of sex steroids. However, small fish such as zebrafish and medaka, which are particularly good model organisms considering the well-developed genetic toolkit and the numerous protocols available to investigate their biology and physiology, raise challenges for applying such protocols due to their small size. Here, we demonstrate the step-by-step procedure of gonadectomy in both males and females followed by blood sampling in a small sized teleost model, the Japanese medaka (*Oryzias latipes*). The use of these procedures combined with the other advantages of using these small teleost models will greatly improve our understanding of feedback mechanisms in the neuroendocrine control of reproduction and tissue plasticity provided by sex steroids in vertebrates.

**SUMMARY:** The article describes a quick protocol to gonadectomize and sample blood from small teleost fish, using medaka (*Oryzias latipes*) as a model, to investigate the role of sex steroids in animal physiology.

## Introduction

In vertebrates, sex steroids, which are mainly produced by the gonads, play important roles in the regulation of the Brain-Pituitary-Gonadal (BPG) axis through various feedback mechanisms (reviewed in ^1-5^). In addition, sex steroids also affect the proliferation and activity of neurons in the brain (reviewed in ^6-8^) and endocrine cells, including gonadotropes, in the pituitary (reviewed in ^9,10^), and thus serve crucial roles in brain and pituitary plasticity.

Despite a relatively good knowledge in mammals, mechanism of BPG axis regulation mediated by sex steroids is far from being understood in non-mammalian species, leading to poor understanding of evolutionary conserved principles (reviewed in ^11^). Besides, there is still a limited number of studies documenting the role of sex steroids on brain and pituitary plasticity, thus raising the need for further investigations of the role and effects of sex steroids on diverse vertebrate species.

Among vertebrates, teleosts have become powerful model animals in addressing numerous biological and physiological questions, including stress response (reviewed in ^12,13^), growth (reviewed in ^14,15^), nutritional physiology (reviewed in ^16,17^) and reproduction (reviewed in ^2^). Teleosts, in which sex steroids are mostly represented by estradiol (E2) in females and 11-ketotestosteron (11-KT) in males ^18,19^, have thus long been reliable experimental models for investigating the general principle of reproduction across species. Also, teleosts possess unique characteristics in their hypothalamus and pituitary, which are sometimes convenient for the elucidation of regulatory mechanisms. For instance, they show direct pituitary innervation of hypophysiotropic neurons in the brain instead of mediating the hypothalamic-hypophyseal portal system (reviewed in ^20,21^) and their gonadotropins (luteinizing and follicle stimulating hormones) are produced in two separate cells in these animals (reviewed in ^22^). Due to the high number of species (nearly 30 000 ^23^) and their high diversity ^24^, these animals offer interesting models to investigate a wide range of biological questions (reviewed in ^25^). Moreover, due to their amenability to both laboratory and field experiments, teleosts offer many advantages compared to other organisms. They are relatively inexpensive to purchase and maintain (reviewed in ^25,26^). In particular, small teleost models, such as zebrafish and medaka, are species with very high fecundity and relatively short life cycle enabling rapid analysis of gene function and disease mechanisms, thus providing even greater advantages in addressing a plethora of biological and physiological questions, considering the numerous well-developed protocols and genetic tools available for these species (reviewed in ^27,28^).

In numerous studies, the removal of gonads (gonadectomy), the main source of sex steroid production, has been used as a method for investigating many physiological questions, including its impact in vertebrate reproductive physiology, including in mammals ^29-31^, birds ^32^ and amphibians ^33^. Meanwhile, blood collection is commonly aimed for quantifying circulating hormone levels, including those of sex steroids ^34-37^. Together, these two techniques have shown their importance in a great number of studies, including the investigation of feedback mechanisms and the effect of sex steroids on BPG axis regulation ^38-40^. In teleosts however, while these techniques are relatively easy to perform in bigger species, such as European sea bass ^41^, coral reef fish ^42^, dogfish ^43^ and catfish ^44,45^, they raise challenges when applied in smaller fishes, such as zebrafish and medaka. Therefore, a clear protocol demonstrating every step of gonadectomy and blood sampling in small teleosts is of importance.

Here, we use Japanese medaka (*Oryzias latipes*) as a model, a small freshwater fish native to East Asia, and similar to zebrafish in many aspects (size, genome sequenced, molecular and genetic tools available). However, medaka has a smaller genome size than that of zebrafish ^46^ and a genetic sex determination system allowing for investigation of sexual differences before second sexual characters or gonads are well developed (reviewed in ^47^).

This paper demonstrates gonadectomy and blood sampling in small teleost models, a technique that takes only 8 minutes in total and that will complete the list of detailed video protocols already existing for this species that included labeling of blood vessels ^48^, patch-clamp on pituitary sections ^49^ and brain neurons ^50^, and primary cell culture ^51^. This technique will allow the research community to investigate and better understand the roles of sex steroids in feedback mechanisms as well as brain and pituitary plasticity in the future.

## Protocol

All experimentations and animal handling were conducted in accordance with the recommendations on the experimental animal welfare at Norwegian University of Life Sciences. Experiments using gonadectomy were approved by the Norwegian Food Safety Authority (FOTS ID 24305).

## 1. Instruments and solutions preparation

1.1. Prepare anesthesia stock solution (0.6% Tricaine):
  1.1.1. Dilute 0.6 g of Tricaine in 100 ml of 10X Phosphate Buffer Saline (PBS).
  1.1.2. Distribute 800 μl of the tricaine stock solution into several 1.5 ml plastic tubes and store at −20 °C until use.
1.2. Prepare recovery water (0.9% NaCl solution) by adding 18 g of NaCl into 2 L of aquarium water. Store the solution at room temperature until use.
1.3. Prepare the incision tools by breaking a razor diagonally to get a sharp point **(Figure 1A)**.
1.4. Prepare blood anti-coagulant solution (0.05 U/μl of sodium heparin) by diluting 25 μl of sodium heparin into 500 μl of 1X PBS. Store the anti-coagulant solution at 4 °C until use.
1.5. Prepare two glass needles from a 90 mm long glass capillary by pulling a glass capillary with a needle puller **(Figure 1B)** following the instructions of the manufacturer.
**NOTE:** The outer diameter of the glass needle is 1 mm, while the inner diameter is 0.6 mm.
1.6. Prepare a 1.5 ml plastic tube lid by cutting the lid and make a hole that fits with the needle outer diameter **(Figure 1C)**. To make the hole, heat one end of the 9-mm glass capillary and stab the heated glass capillary through the lid. Alternatively, use a needle to stab through the lid until the diameter of the hole fits with the 9-mm glass capillary.

**Fig 1.**
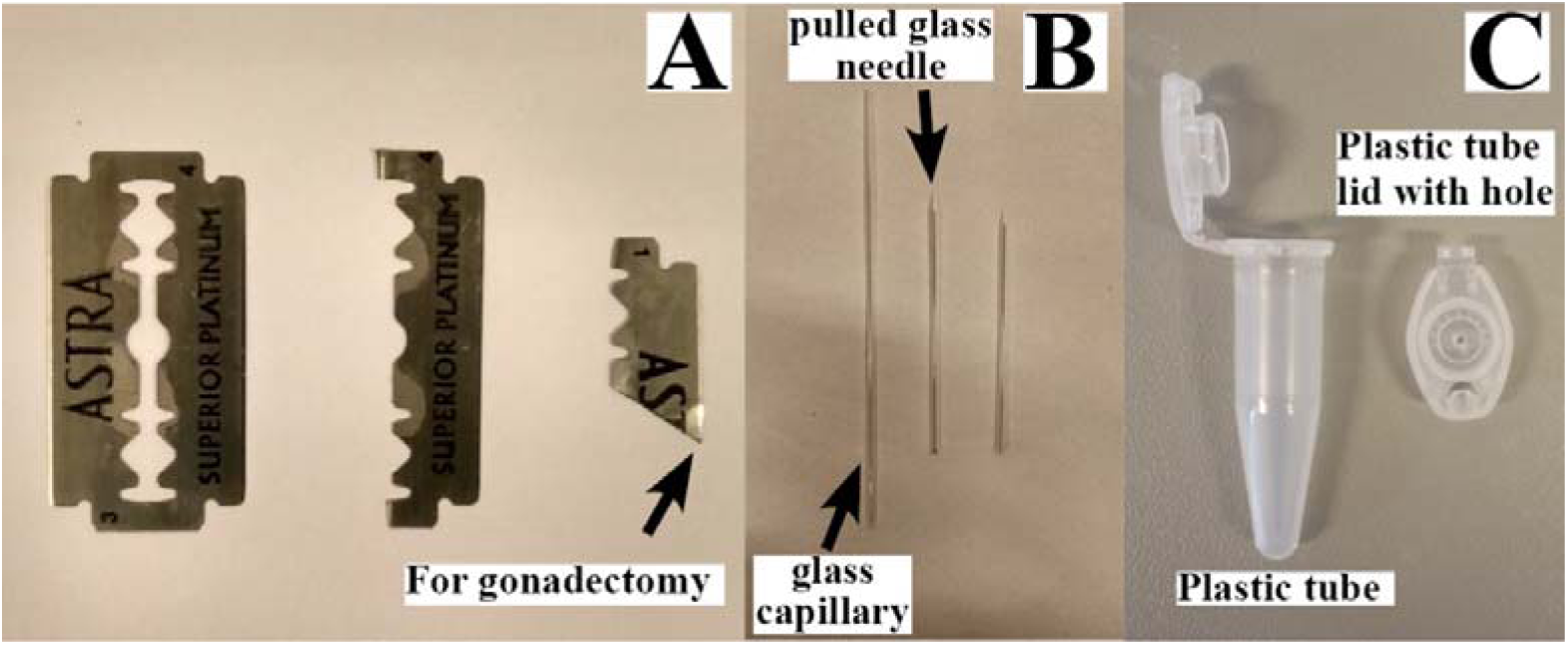
Razor blade for gonadectomy (A), glass needle for blood extraction (B), and plastic tube together with a lid with a hole for blood collection (C).

## 2. Gonadectomy procedure

2.1. Prepare 0.02% of anesthesia solution (MS-222) by diluting one tube of Tricaine stock (0.6%) in 30 ml of recovery water.
**NOTE:** Depending on the size of the fish, additional Tricaine stock can be added, but final concentration of 0.02 % usually works well.
In our experience, most cases that fish did not recover after surgery were due to over exposure to high concentration of anesthesia, probably not due to the mistake in surgery.
2.2. Prepare dissection tools including one ultra-fine and two fine forceps (one with relatively wide tip), small scissors, nylon thread and razor as described in **step 1**.**3**.
2.3. Anesthetize the fish by putting it into the 0.02% anesthesia solution, and make sure that the fish is anesthetized enough to be operated.
**NOTE:** To ensure that the fish is fully anesthetized, the fish body can be pinched gently using forceps. If the fish does not react, the gonadectomy can be started.
2.4. Place the anesthetized fish under a dissection microscope.
2.5. Ovariectomy in females
  2.5.1. Remove oviposited eggs (eggs hanging outside the female body) if any, and scrap the scales in the incision area **(Figure 2A)**.
  2.5.2. Incise gently the incision area between the ribs **(Figure 2A)** using the razor blade, pinch gently the fish abdomen while taking out the ovary little by little using fine forceps with wide tip.
  2.5.3. Cut the end of the ovary using fine forceps and put aside the ovary **(Figure 2B)**.
**NOTE:** It is important to take care not to break the ovarian sac as possible. In case of breaking the ovarian sac, it is important to remove as completely as possible without leaving even some non-ovulated eggs.
2.6. Orchidectomy in males
  2.6.1. After making an incision with the razor blade, incise gently between the ribs **(Figure 2A)** and open up the incision slowly using fine forceps.
  2.6.2. Grab the testis gently using the fine forceps and take out the testes slowly. Afterwards, cut the end of the testis to remove testis completely **(Figure 2B)**.
**NOTE:** For male orchidectomy, all preparations are similar to in females until the incision part. Incision area of males should be more dorsal side of the abdomen **(Figure 2A)**. When grabbing the testes, sometimes we obtain only the fat resembling the testes. However, after restoring the fat, it is possible to try to find the testes again. **(Figure 2B)**.
**NOTE:** For both males and females, it is important to minimize the incision size in the abdomen to prevent excessive damage that can lead to mortality. Sometimes the intestines may also appear through the incision along with the gonads, so make sure they are properly returned inside the incision before closure. It is important to understand where ovaries or testes are localized in medaka abdomen by dissection.
2.7. Suture the incision similarly in males and females **(Figure 3)**.
  2.7.1. Place the nylon thread beside the incision area, inject the right side of incision part from inner body cavity using ultra-fine forceps to take the thread in with the help of fine forceps **(Figure 3;1-2)**.
  2.7.2. Inject the left side of incision part from outer body cavity to take out the thread **(Figure 3;3-4)**.
  2.7.3. Close the incision opening and make two knots and cut the excessive thread **(Figure 3;4-6)**.
  2.7.4. Put the fish directly into the recovery water.
**NOTE:** The suture should be adequately tight, and the remaining thread on the fish should be long enough to prevent the disattachment of the suture.

**Fig 2.**
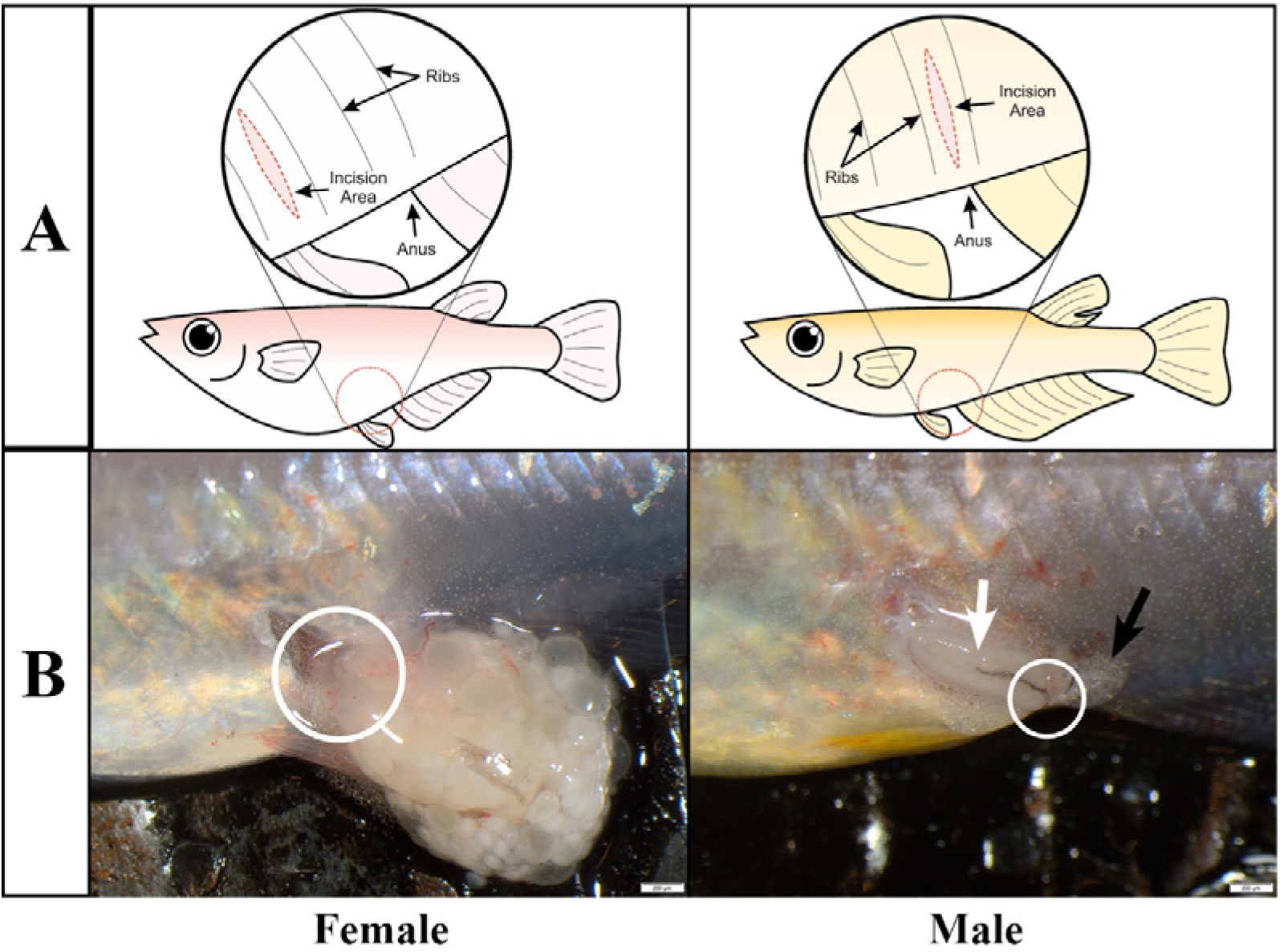
Location of the incision area. A) Drawing of the incision area located between the ribs in females (left panel) and males (right panel); B) gonad removal in females (left panel) and males (right panel), white circles showing the joint part, white arrow showing the testis and black arrow showing the fat.

**Figure 3.**
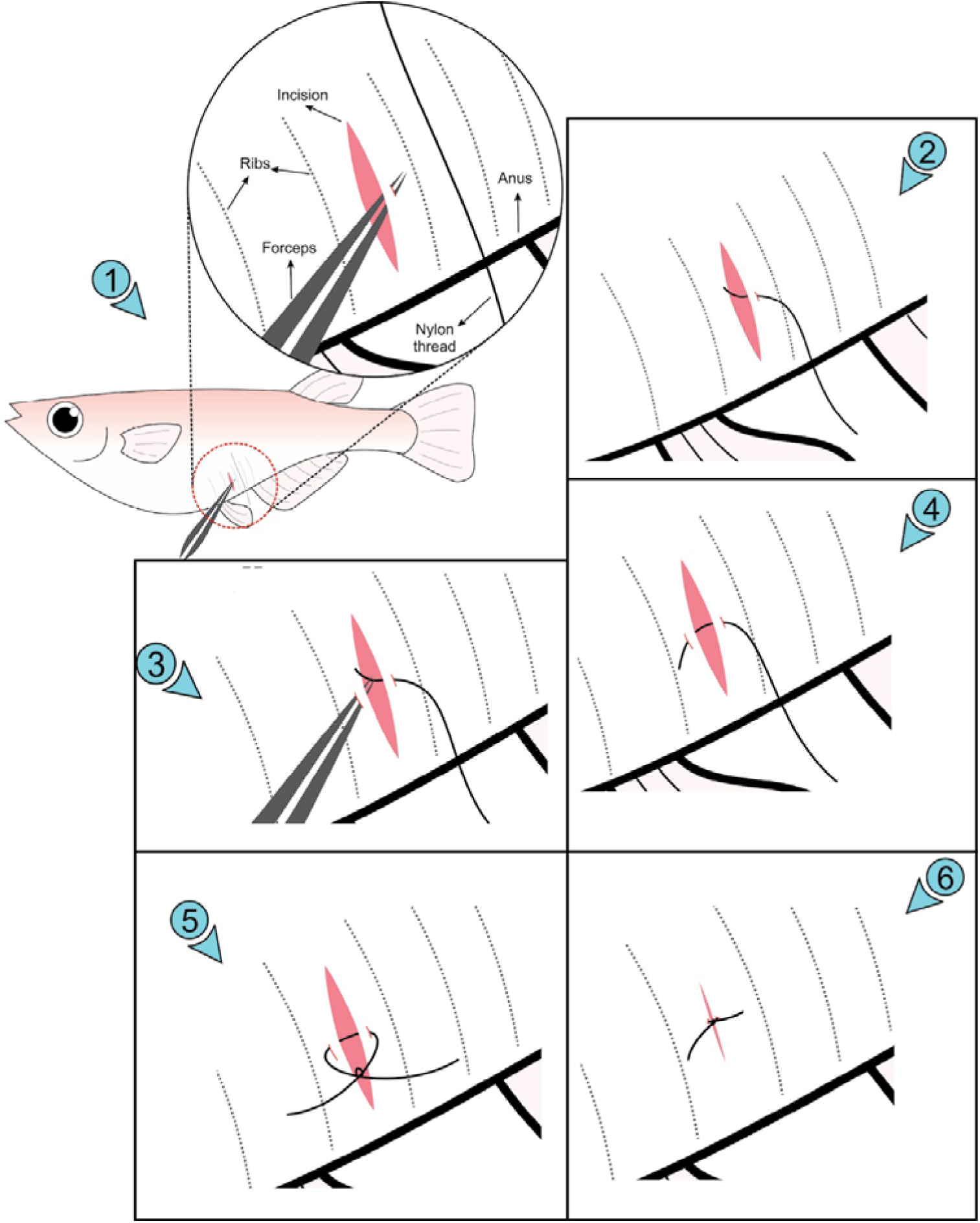
The procedure of suture. 1) a hole is made on the right side of the incision using fine forceps. 2) the nylon thread is passed through the skin using the hole made in 1. 3) a hole is made in the left side of the incision. 4) the nylon thread is passed through the hole made in 3. 5) an overhand knot is made twice to close the incision. 6) excess thread is cut.

## 3. Blood sampling procedure

3.1. Prepare the tools including glass needle, silicone capillary, a plastic tube with a hole, an empty 1.5 ml plastic tube, a 1.5 ml plastic tube containing 1X PBS, mini centrifuge, and tape.
3.2. Anesthetize the fish using 0.02% MS-222 solution as described in **step 2**.**1**, and place the fish under a dissection microscope in a vertical position **(Figure 4A)**.
**NOTE:** It is highly recommended to place the fish on a bright surface to ease visualization of the caudal puncture vein.
3.3. Install the blood drawer by attaching a glass needle to the silicone tubing **(Figure 4B)**. Break the tip of the needle with forceps with wide tip, and coat the anti-coagulant inside a needle by suctioning and blowing.
**NOTE:** Make sure that the opening of the needle tip is sufficiently large to allow drawing the blood.
3.4. Direct the needle toward the peduncle area of the fish, aim at the caudal peduncle vein **(Figure 5A)** and draw the blood using mouth until at least one fourth the total volume of the needle is filled **(Figure 5B)**.
**NOTE:** It is important to stop suctioning when removing the needle from fish body.
3.5. Release the needle and put a piece of tape on it. Place the lid with a hole on a reservoir tube and put the needle inside the tube through the hole with the needle tip on the outside **(Figure 5C)**.
3.6. Spin down the blood to collect the blood in the tube
3.7. Proceed directly to downstream applications, or store the blood at −20 °C until use.
**NOTE:** The blood analysis is dependent on what outcome is required. For sex steroid analysis such as E2 or 11KT, 1 ul of collected blood is adequate for the analysis. The blood can be diluted in 1X PBS and extract steroids with diethyl ether or dichloromethane if necessary. In many previous studies, clot was removed, however, as the volume of blood is so small that we can ignore the effects in most cases.

**Fig 4.**
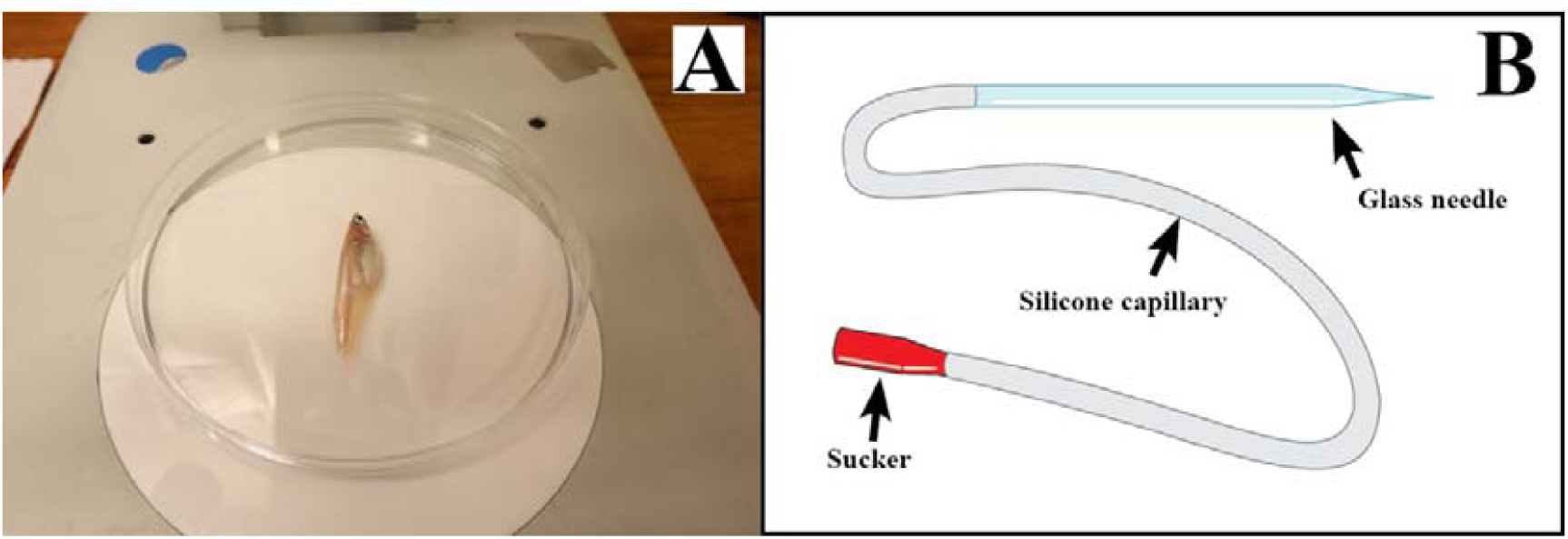
Fish position during blood sampling (A), the installation of glass needle with the silicone capillary (B).

**Fig 5.**
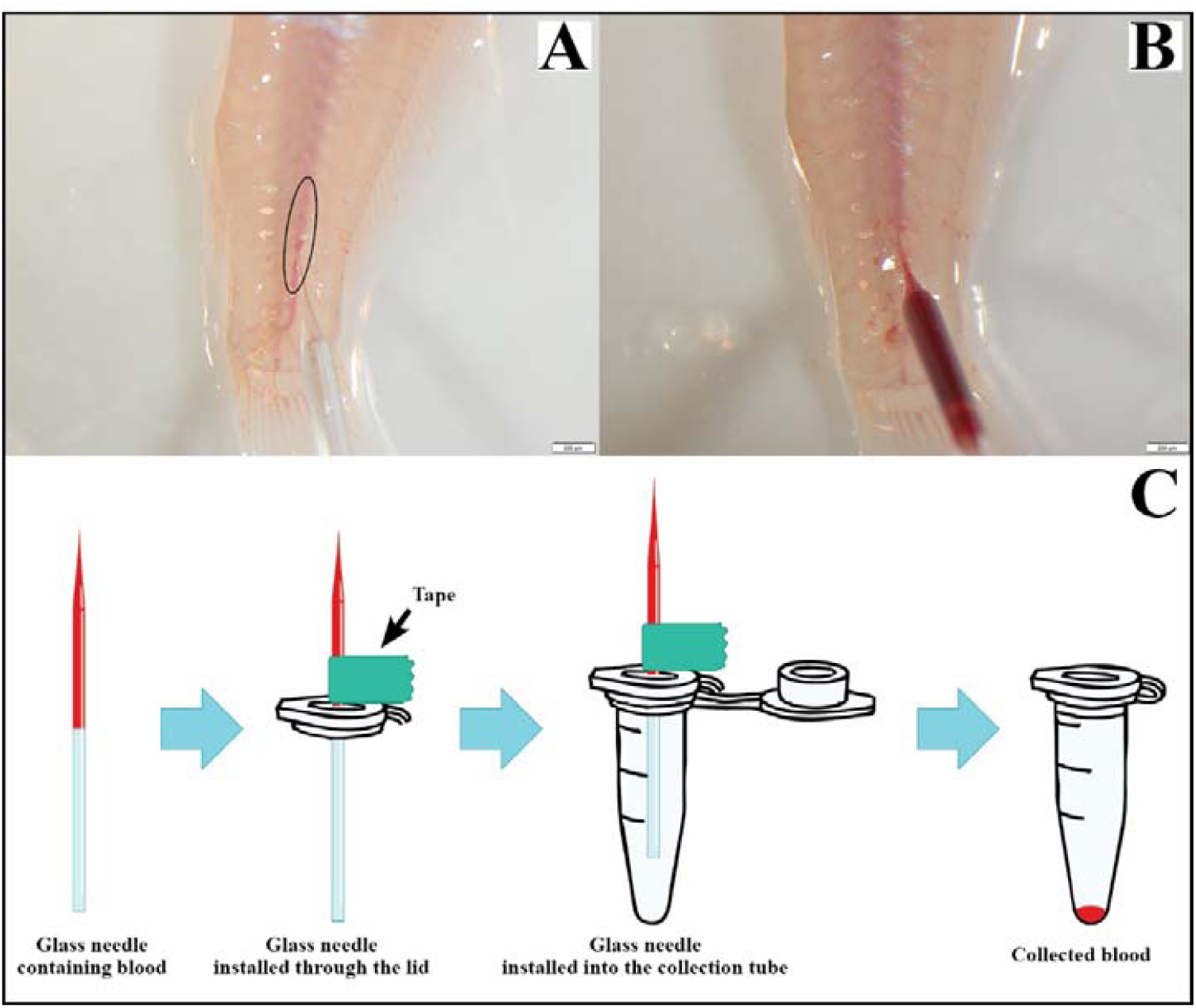
The suction area of blood sampling (A), drawn blood (B) and blood collection steps (C).

## Representative Results

This protocol describes every step for performing gonadectomy and blood sampling in small sized teleosts, using the Japanese medaka as a model. The survival rate of the fish after ovariectomy (OVX) in females is 100% (10 out of 10 fish) while 94% (17 out of 18 fish) of the males survived after orchidectomy. Meanwhile, after blood sampling procedure was performed, all (38 fish) fish survived.

Sham-operated females show oviposition **(Figure 6A)** and all the eggs are fertilized and allow for embryonic development **(Figure 6B)**. Sham operated males are also able to fertilize eggs after only a couple of weeks. Similarly, partly-gonadectomized females reared with partly-gonadectomized males also show oviposition and showed 100% of fertilized eggs after 2 months. In contrast, no oviposition in females or fertilization by males could be observed in fully gonadectomized fish, even after 4 months.

**Fig 6.**
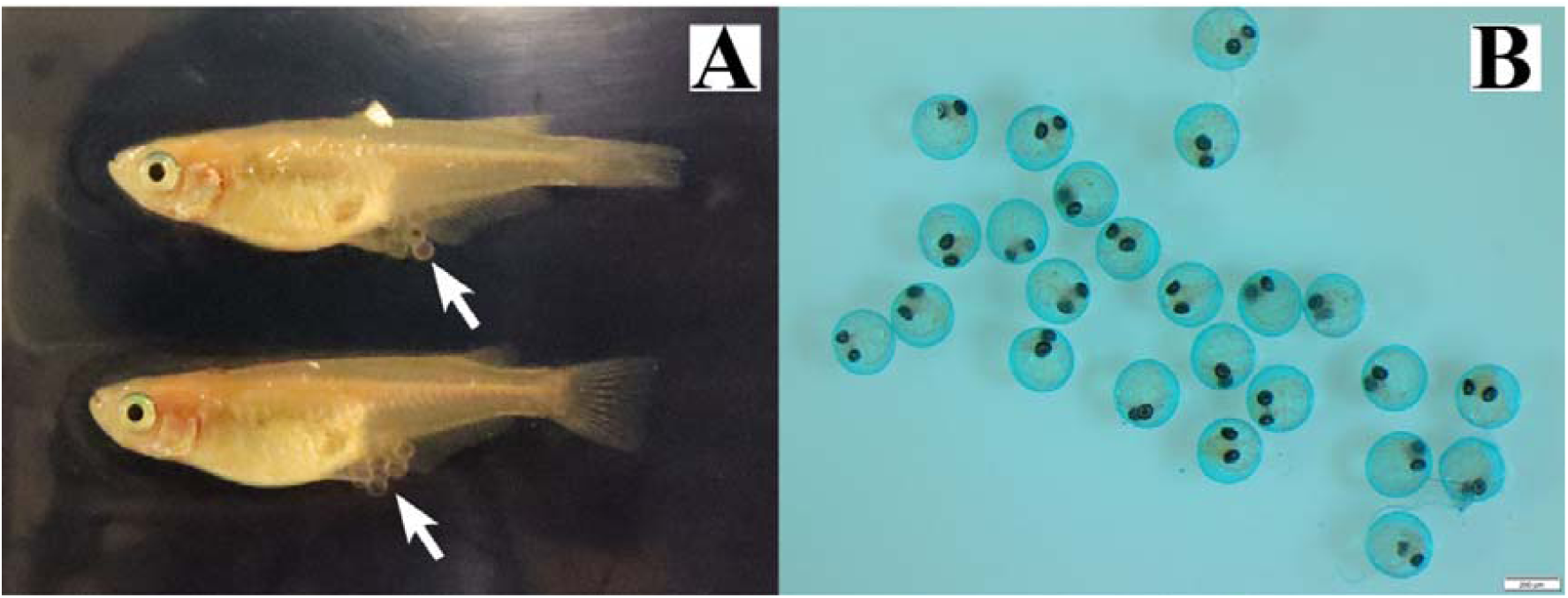
Sham-operated fish shows spawning. Oviposition of eggs (A) and fertilized eggs (B).

When performed correctly, the body shape of the fish slightly changes (Figure 7A). If performed correctly, no piece of gonad should remain after the gonadectomy procedure when dissected (Figure 7B).

**Fig 7.**
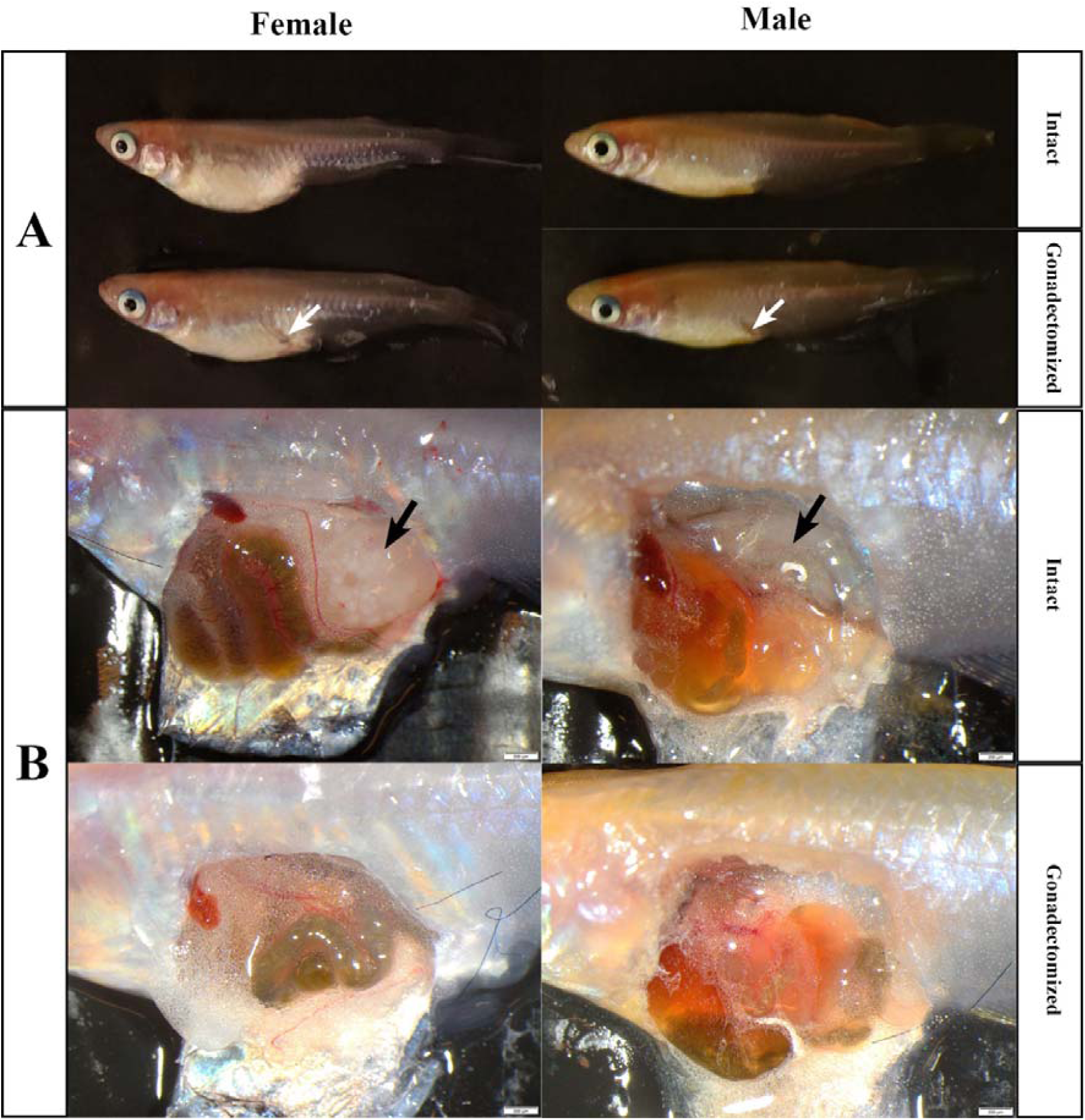
Morphological (A) and anatomical (B) appearance of intact and gonadectomized fish

Four weeks post-gonadectomy, the incision and suture completely disappeared **(Figure 8)**, and after 4 months, all gonadectomized fish still showed healthy phenotype, and no gonadal tissue could be found.

**Fig 8.**
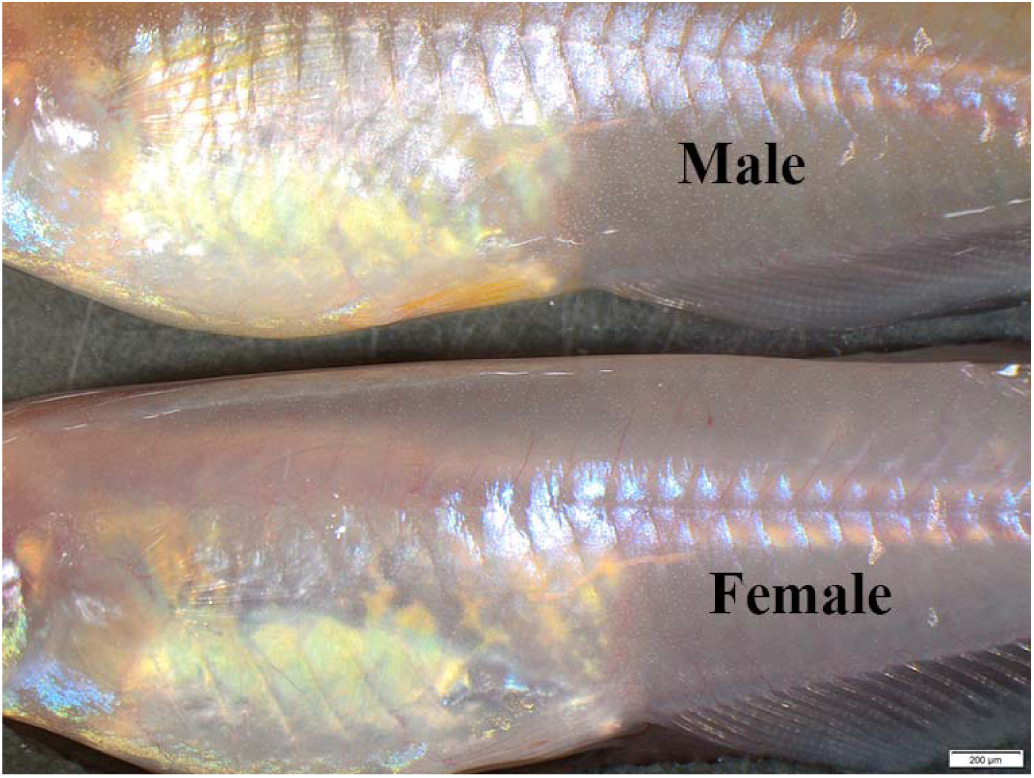
The incision and suture after 4 weeks.

E2 and 11-KT blood concentrations measured with ELISA following the manufacturer’s instructions revealed that E2 levels in OVX females (0,36 ± 0,2 ng/ml) are significantly lower than in sham-operated females (4,15 ± 0,5 ng/ml) 24 hours after surgery **(Figure 9A)**. Likewise, 11-KT concentrations in orchidectomized males (0,4 ± 0,2 ng/ml) are also significantly lower than in sham-operated males (10.38 ± 1.32 ng/ml) 24 hours after surgery **(Figure 9B)**. There is no statistical difference in blood levels of E2 and 11KT in gonadectomized fish after 4 months compared to the levels of those after 24 hours **(Figure 9A-B)**.

**Fig 9.**
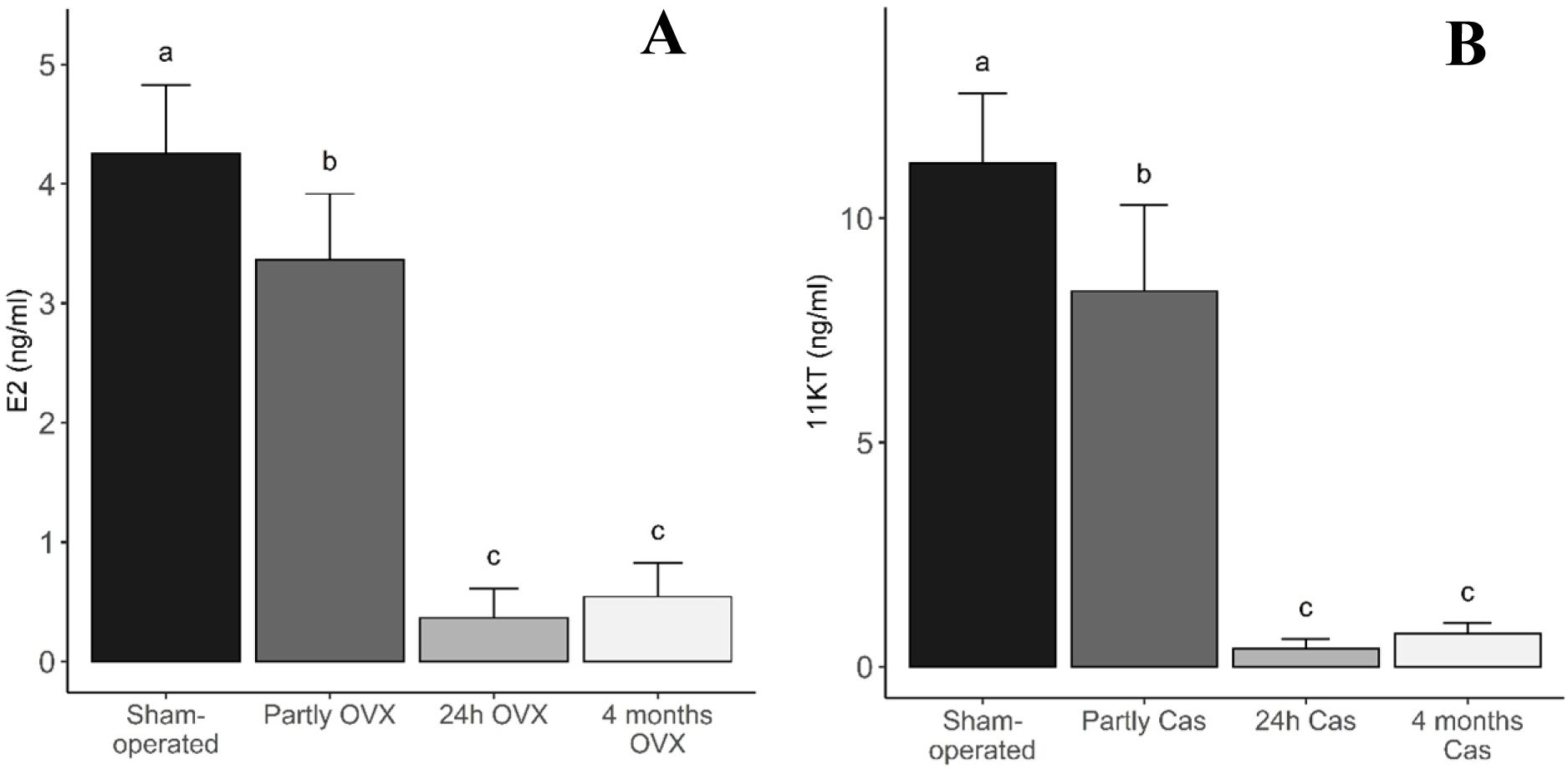
Blood levels of E2 in female (A) and 11KT in male (B) medaka, 24 hours after sham operation (control), partly gonadectomy or gonadectomy, and 4 months after gonadectomy (OVX, ovariectomy in females; Cas, castrated in males) (Data in the graph are provided as mean + SD; n = 5). Different letters (a-c) show statistically significant differences.

In contrast to fully OVX females, partly OVX fish, where only 1/3 to 1/2 of the gonad was removed, showed no difference in E2 levels (3,37 ± 0,6 ng/ml) compared to sham-operated fish **(Figure 9A)**. However, in males there is a difference observed between 11KT levels of sham-operated fish and partly orchidectomized fish (8.37 ± 1.92 ng/ml) **(Figure 9B)**. However, partly orchidectomized males, where only 1/3 to 1/2 of the gonad was removed, showed significantly higher levels of 11KT compared to in fully orchidectomized fish.

## Discussion

As reported in previous literature, gonadectomy and blood sampling have long been used in other model species to investigate questions related to the role of sex steroids in regulation of the BPG axis. However, these techniques seem to be amenable only for bigger animals. Considering the small size of the most used teleost models, we hereby describe detailed protocols for gonadectomy and blood sampling that are feasible for these small teleost models.

The fact that the survival rate of gonadectomized fish reached almost 100% indicates that the gonadectomy procedure is feasible to be applied on small fish. Similarly, the procedure of blood sampling does not affect the survivability of the fish as shown by the 100% survival after undergoing this procedure. In addition, sham-operated females reared together with sham-operated males show oviposition and 100% fertilized eggs, indicating that the incision and suture procedure do not affect the reproduction of the fish. In other words, they were healthy enough to spawn. Meanwhile, as shown in **Figure 7**, the incision and suture mark on the fish completely disappeared 4 weeks post-gonadectomy and the fish are still alive and look healthy 4 months after surgery, indicating that the operation procedure is safe for the fish for long term purpose gonadectomy and does not affect the life span of the fish. In addition, after 4 months no gonads are observed. This is confirmed by the low levels of E2 and 11KT which are still similar to that of those found in gonadectomized fish after 24 hours.

Partly gonadectomized fish showed comparable concentrations of sex steroids to sham-operated fish, and as a consequence, resulted in oviposition in the females and fertilization of eggs. These results suggest that the procedure of gonadectomy should be performed with high precision, meaning that the ovary or testes should be completely removed. Furthermore, since this procedure does not rely on Fish Anesthesia Delivery System (FADS) as demonstrated in ^52^, the gonadectomy should be carried out as quickly as possible to prevent mortality during surgery. Indeed, the use of FADS enables us to maintain the rhythm of operation since this tool allows continuous anesthetic condition to the fish despite being exposed to the air. Nonetheless, due to its lower feasibility in smaller teleosts, the use of FADS cannot be performed with these sized fish. Many factors can affect the success rate of the procedure, including anesthesia period, the wideness of incision, the accuracy and tidiness of the suture and fish handling during the procedure. Since the protocol relies so much on the quick and clean procedure, some training is highly recommended until reaching high success rate, indicated by high survival rate of the fish after gonadectomy as well as complete removal of the gonads (see the difference of morphological and anatomical appearance of the fish before and after successful gonadectomy in **Figure 7**). Another important point is that one should prepare healthy fish by maintaining the fish optimally prior to performing the protocol.

With respect to blood sampling procedure, the steroid extraction is generally performed using diethyl ether or dichloromethane. Meanwhile, the evaluation of sex steroid concentrations is commonly carried out by using Enzyme-linked Immunosorbent Assay (ELISA) kit, and there have been many ELISA kits commercially available for different types of sex steroids. Due to the low amount of blood collected during blood sampling, the assays performed to evaluate sex steroid concentrations in small teleosts is not aimed for serum, but whole blood. This might influence the results obtained from the assays. A previous study ^53^ suggested that the quantification of sex steroids using whole blood can slightly differ from that of serum. Therefore, the comparison between plasma and blood concentrations of sex steroids should be investigated for each assay in order to determine whether measured concentrations from the assay should be re-calculated to get comparable results as from serum.

As documented in previous studies with different animal models, the protocol described here will allow us to investigate questions related to reproductive physiology using small teleosts as model. In fact, these techniques have already contributed to answer questions concerning the regulation of the BPG axis and its feedback mechanisms, such as the involvement of *kiss1* (kisspeptin gene type 1) expressing neurons in positive feedback loops ^54^, estrogen-mediated regulation of *kiss1* expressing neurons in nucleus ventralis tuberis (NVT), and *kiss2* (kisspeptin gene type 2) expressing neurons in preoptic area (POA) ^55,56^, the expression profile of *fshb* (follicle-stimulating hormone beta sub-unit gene) in *esr2a* (estrogen receptor gene) knock out (KO) fish ^57^ as well as the profile of circadian rhythm of E2 in female fish ^53^. Furthermore, since previous studies demonstrated that sex steroids also affect the proliferation of gonadotropic cells in the pituitary of teleosts ^58,59^, it would be intriguing to investigate the effects of sex steroid clearance after gonadectomy on pituitary plasticity. Besides, due to the fact that the protocol can also be applied for blood glucose measurements as demonstrated in zebrafish ^60^ and medaka ^61^, it may also be expanded to address research questions in other fields of physiology.

Finally, the protocols described here are intended and optimized for adult medaka, and the outcomes due to different size of fish and materials used during the procedures may vary. Also, as medaka left and right ovaries/ testes are fused, which might provide an important advantage for gonadectomy, this protocol might need few small adaptations before to be used in other species where this is not the case such as in zebrafish. Thus, an optimization according to the choice of laboratory equipment and fish size should be taken into account before testing these protocols.

## Discolsures

The authors have nothing to disclose.

## Acknowledgements

The authors thank Ms Lourdes Carreon G Tan for her assistance in the fish husbandry. This work was funded by NMBU, Grants-in-Aid from Japan Society for the Promotion of Science (JSPS) (Grant number 18H04881 and 18K19323), and grant for Basic Science Research Projects from Sumitomo Foundation to S.K.

## Notes

### Competing Interest Statement

The authors have declared no competing interest.

